# Definition, modeling and detection of saccades in the face of post-saccadic oscillations

**DOI:** 10.1101/2021.03.24.436800

**Authors:** Richard Schweitzer, Martin Rolfs

## Abstract

When analyzing eye tracking data, one of the central tasks is the detection of saccades. Although many automatic saccade detection algorithms exist, the field still debates how to deal with brief periods of instability around saccade offset, so-called post-saccadic oscillations (PSOs), which are especially prominent in today’s widely used video-based eye tracking techniques. There is good evidence that PSOs are caused by inertial forces that act on the elastic components of the eye, such as the iris or the lens. As this relative movement can greatly distort estimates of saccade metrics, especially saccade duration and peak velocity, video-based eye tracking has recurrently been considered unsuitable for measuring saccade kinematics. In this chapter, we review recent biophysical models that describe the relationship between pupil motion and eyeball motion. We found that these models were well capable of accurately reproducing saccade trajectories and implemented a framework for the simulation of saccades, PSOs, and fixations, which can be used – just like datasets hand-labelled by human experts – to evaluate detection algorithms and train statistical models. Moreover, as only pupil and corneal-reflection signals are observable in video-based eye tracking, one may also be able to use these models to predict the unobservable motion of the eyeball. Testing these predictions by analyzing saccade data that was registered with video-based and search-coil eye tracking techniques revealed strong relationships between the two types of measurements, especially when saccade offset is defined as the onset of the PSO. To enable eye tracking researchers to make use of this definition, we present and evaluate two novel algorithms – one based on eye-movement direction inversion, one based on linear classifiers previously trained on simulation data. These algorithms allow for the detection of PSO onset with high fidelity. Even though PSOs may still pose problems for a range of eye tracking applications, the techniques described here may help to alleviate these.

## 1 Introduction

Rapid, step-like eye movements, so-called saccades, are the fastest and most frequent of human movements. As they reorient the fovea – the retina’s area of sharpest vision – towards objects of interest and produce transients in the input, saccades are crucial for the active perception of the visual world. For more than a century, researchers have measured saccades and their role in visual perception using various eye tracking techniques, such as visual inspection, mechanical recordings, tracking of mirror-reflected beams of light, electrooculography, and even the apparent movement of afterimages [1]. Today, the most common applications use the method of video-based eye tracking. Using infrared or near-infrared light sources, video-based systems track the center of the illuminated pupil, as well as position of the first Purkinje image (PI), the corneal reflection, to estimate gaze position.

Due to the high speeds of saccades, which, at large amplitudes, routinely reach peak velocities of up to 700 degrees of visual angle per second (dva/s) [2], inertial forces act on the eye. These forces act not only on photoreceptors on the retina, which tilt away from the direction of rotation [3], but also on the lens and the iris, whose relative motion cause so-called post-saccadic oscillations (PSOs), widely also referred to as dynamic overshoots [4]. These can be observed in eye tracking data as brief periods of instability, or even ringing, around the end of a saccade (**Figure 1a,b**), often with durations of ~25 ms [5]. Alternatively, if the eye is filmed by a high-speed camera, a PSO can be identified as a brief deformation or “wobbling” of the iris following a saccade. In the case of video-based eye tracking, there is good evidence that it is specifically the motion of the pupil within the iris that causes prominent PSOs [6–8], whereas in Dual Purkinje Image (DPI) eye tracking systems, they are related to the misalignment of cornea (1st and 2nd PIs) and lens (3rd and 4th PIs) [9–11]. To elucidate, a short slow-motion video, which we recorded with a Phantom high speed camera at 7300 fps and that can be viewed at https://osf.io/23798/, shows not only post-saccadic movement of the pupil border relative to the prominent corneal reflection, but also a weaker light spot within the pupil, most likely a reflection from the lens, with striking lag and oscillation. Yet, it has been suggested that at least a small proportion of PSOs has its origin in oculomotor control signals, specifically in the temporal coordination of agonist and antagonist motoneural activity [4, 12, 13]. There is evidence for this view, as small PSOs could also be shown when measuring saccades with scleral search coils, attached to the eye’s cornea and thus directly measuring the rotation of the eyeball [14–17]. However, these PSOs – provided that they were not caused by low contact-lens slippage – occurred mainly for very short saccades or microsaccades, had significantly lower incidence and smaller amplitudes than in video-based systems, and contained not more than one phase [18, 19].

**Fig. 1:**
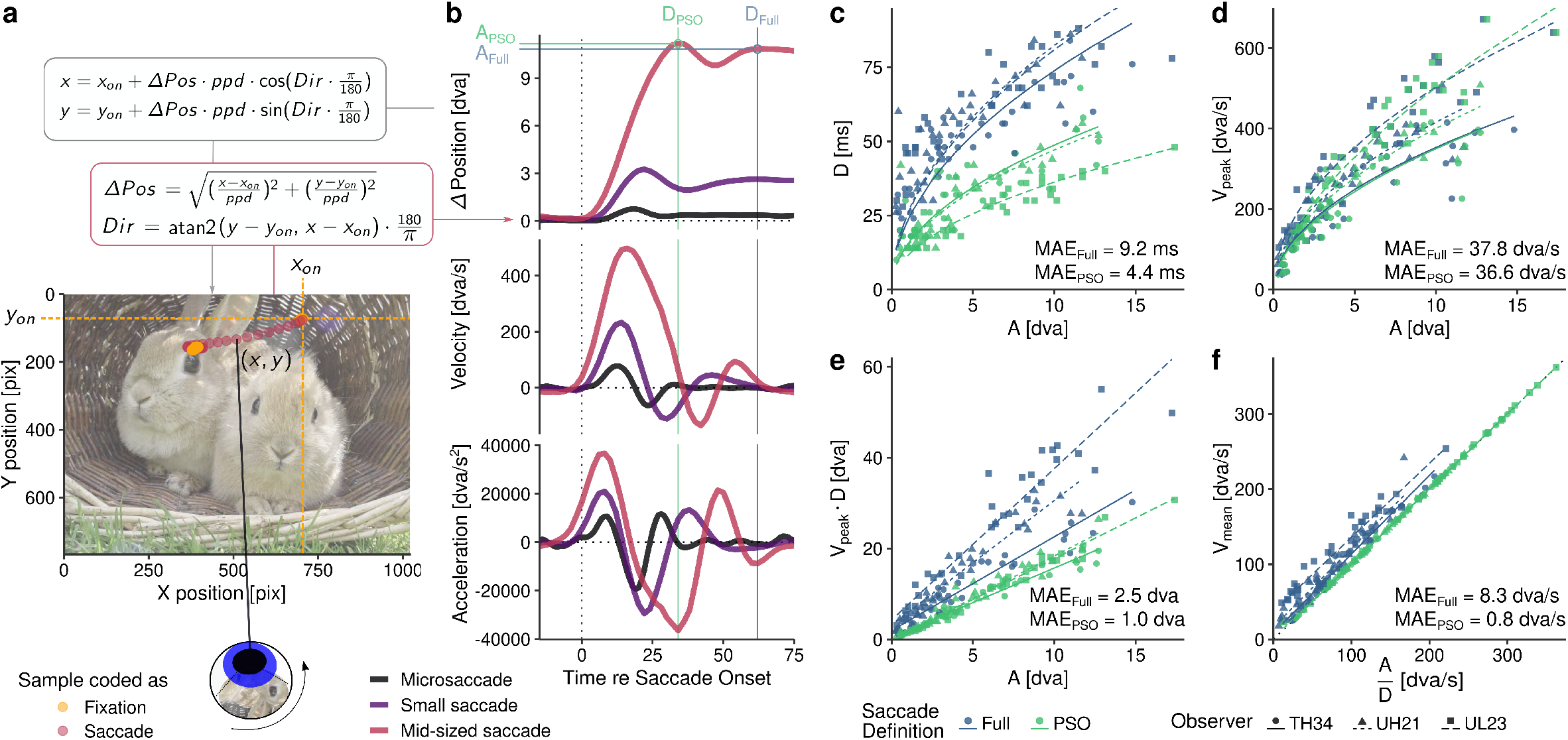
Illustration of different saccade definitions. **a** How to transform a two-dimensional gaze position (*x*, *y*) signal to position change (Δ*Pos*) and direction (*Dir*) over time to visualize post-saccadic oscillations. The data [taken from a handlabeled data set, 21–23] shows a leftward saccade that is initiated at fixation position (*x*_*on*_, *y*_*on*_). The constant *ppd* (pixels per degree) transforms eye position in pixel coordinates into degrees of visual angle. **b** Position, velocity, and acceleration profiles of three saccades of different sizes, of which one is a microsaccade [from a hand-labeled data set, 24]. Saccade amplitudes and durations may be computed based on different offset definitions, namely the onset or offset of the post-saccadic oscillation (*D*_*PSO*_ or *D*_*Full*_), respectively. **c-f** Saccadic main sequence relationships for three observers [again from 21–23], depending on the PSO (green) and Full (blue) saccade offset definitions. Amplitude-duration relationships were fitted with a square-root model (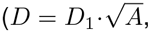, [25]) and amplitude-peak velocity relationships were fitted by a exponential model (*V*_*peak*_ = *V*_1_ · *A*^*P*^, [2]), using the mixed-effects framework provided by the *nlme* R package [26]. Other relationships are approximated by linear mixed-effects models fitted with the *lme4* R package [27]. MAE indicates the mean absolute errors of each set of fits, separately for each saccade offset definition.

In the context of video-based eye tracking, the notion that PSO amplitude should decrease with saccade amplitude is captured in the “gentle braking” hypothesis: Short saccades must decelerate more abruptly than large saccades, causing the elastic components of the eye to oscillate to a larger extent [6, 20]. A rule of thumb is that the larger the anatomical distance between an elastic component and the point where the extraocular muscles exert force on the eyeball, the more prominent is the PSO [20]. Indeed, PSO amplitudes are maximal when measured based on the reflections from the lens (4th PI), considerably smaller when measured based on pupil position, and basically absent in the corneal reflection (1st PI) [11].

Why should the eye tracking researcher care about post-saccadic oscillations? Clearly, the question how we treat PSOs is at the core of the (more conceptual) question how we define saccade offset. On the one hand, one may define saccade offset as the point when the elastic components of the eye are at “rest” (even though due to fixational eye movements and tremor the eye is never really static [28, 29]), that is, after the PSO (*Full* definition in **Figure 1**). This is a reasonable definition in the light of the evidence that suggests that the movement of the lens can have manifest perceptual consequences [11, 30]. On the other hand, as several co-registration studies have shown that PSOs may by far outlast the rotation of the eyeball [10, 18, 31], this criterion may lead to gross overestimations of saccade duration. In fact, as shown in **Figure 1b**, PSOs may almost be as long as the main component of the saccade. In cases in which a more conservative criterion is needed, for instance when a visual stimulus manipulation must be concluded while a saccade is still ongoing [32], it may be useful to define the prominent peak of the first oscillation phase (i.e., the onset of the PSO) as the saccade’s offset (*PSO* definition in **Figure 1**). As we will later also show, there is good evidence that “the peak of each overshoot coincides approximately with the time of the conclusion of the coil-measured saccade” [10, p. 532], suggesting that onset of the PSO might be a better predictor for the end of the physical rotation of the eyeball than PSO offset.

With video-based eye tracking data alone, it is hard to evaluate such assumptions as a ground truth is usually unavailable. Yet, researchers have the possibility to assess the plausibility of their saccade definitions by examining the so-called main sequence, which describes stereotypical relationships between saccade amplitude, saccadic peak velocity, and saccade duration [12]. Main sequence relationships can be fitted with a number of models whose goodness of fit indicates how well one’s definition of a saccade adheres to general principles of oculomotor function [33]. Figure 1c indicates that adopting a *Full* saccade definition essentially leads to almost a doubling of saccade duration relative to the *PSO*-defined saccade duration, which is especially drastic at short saccade amplitudes and leads to a strong reduction of the goodness of fits. The same applies to the relationship of saccade amplitude with the product of peak velocity and duration (**Figure 1e**), which should be especially tight as it includes the proportional relationship of peak and mean velocity of saccades [34]. Indeed, for *PSO*-defined saccades, but not for *Full*-defined saccades, mean velocity (computed based on average sample-to-sample velocity) directly translates to the quotient of saccade amplitude and duration (**Figure 1f**), as suggested by many previous investigators [2]. Including the PSO duration to the saccade offset definition may thus disrupt these relationships. The perhaps most frequently described relationship of saccade amplitude and peak velocity, however, remains largely unaltered by saccade offset definitions (**Figure 1d**). This is expected as a saccade’s peak velocity is not reached during the PSO component. Defining the peak of the post-saccadic overshoot as saccade offset necessarily leads to an overestimation of saccade amplitude, but as saccadic peak velocity measured by video-based systems is also overestimated compared to those measured by scleral search coils [6, 10, 18, 19, 35], the relationship can still hold. Yet, saccade amplitudes, if defined as the distance between fixation positions and thus not including PSO-related overshoots, are extremely similar in both video-based and search-coil systems [31, 36]. Thus, not only for compatibility with search-coil data, but also because oscillations imply a repetitive movement around a point of equilibrium, it is more reasonable to use the *Full* definition of saccade amplitude.

The definition of saccade offset determines a vast number of metrics frequently computed by eye tracking researchers, such as fixation duration or (secondary) saccade latency, and may critically affect various applications, ranging from gaze-contingent displays to clinical diagnostics. In fact, some authors went as far as to declare video-based eye tracking unsuitable for studying the dynamics of saccades [6]. This book chapter shall thus discuss potential ways to alleviate these issues. In **section 2.1**, we will describe physiologically plausible models that allow not only for the close approximation of profiles of saccades with post-saccadic oscillations, but also for the extraction of the underlying profiles of eyeball rotation [37, 38]. These models’ predictions will be validated in **section 2.2** on the basis of co-registered video-based and scleral-coil eye tracking data [39]. We will also present simple statistical heuristics that show the relationship between the two types of measurements that quantify motion trajectories of the two different anatomical structures measured by these two eye tracking techniques. Emphasizing the usefulness of detecting the onset of PSOs, we will finally present and evaluate two methods for saccade and PSO detection, namely velocity- and direction-based extensions to the widely used Engbert-Kliegl algorithm for microsaccade detection [40, 41] (**section 2.3**), as well as a more sophisticated approach applying linear classifiers trained on simulated saccade data with model-generated labels (**section 2.4**).

## 2 Methods

### 2.1 Modeling saccade trajectories

Saccades usually follow prototypical position profiles that can be nicely visualized when transforming two-dimensional gaze position data to position change over time (**Figure 1a**), thereby normalizing for different saccade directions. Several models to approximate these profiles have been proposed (see **Figure 2a** for a non-exhaustive list), ranging from cumulative distribution functions to dedicated parametric models [42, 43]. These models can easily be fitted using any nonlinear least-squares method, yet they often do not account for PSOs. In fact, PSOs may even distort these models’ estimates of saccade amplitude. An exception is a recent model (**Figure 2a**, right panel) that assumes a harmonic oscillator driven by a step function. This model is capable of closely approximating the shape of both saccade and post-saccadic oscillation [44] using only four parameters: amplitude (*a*), onset delay (*t*_*0*_), duration (*b*), and frequency (*ω*).

**Fig. 2:**
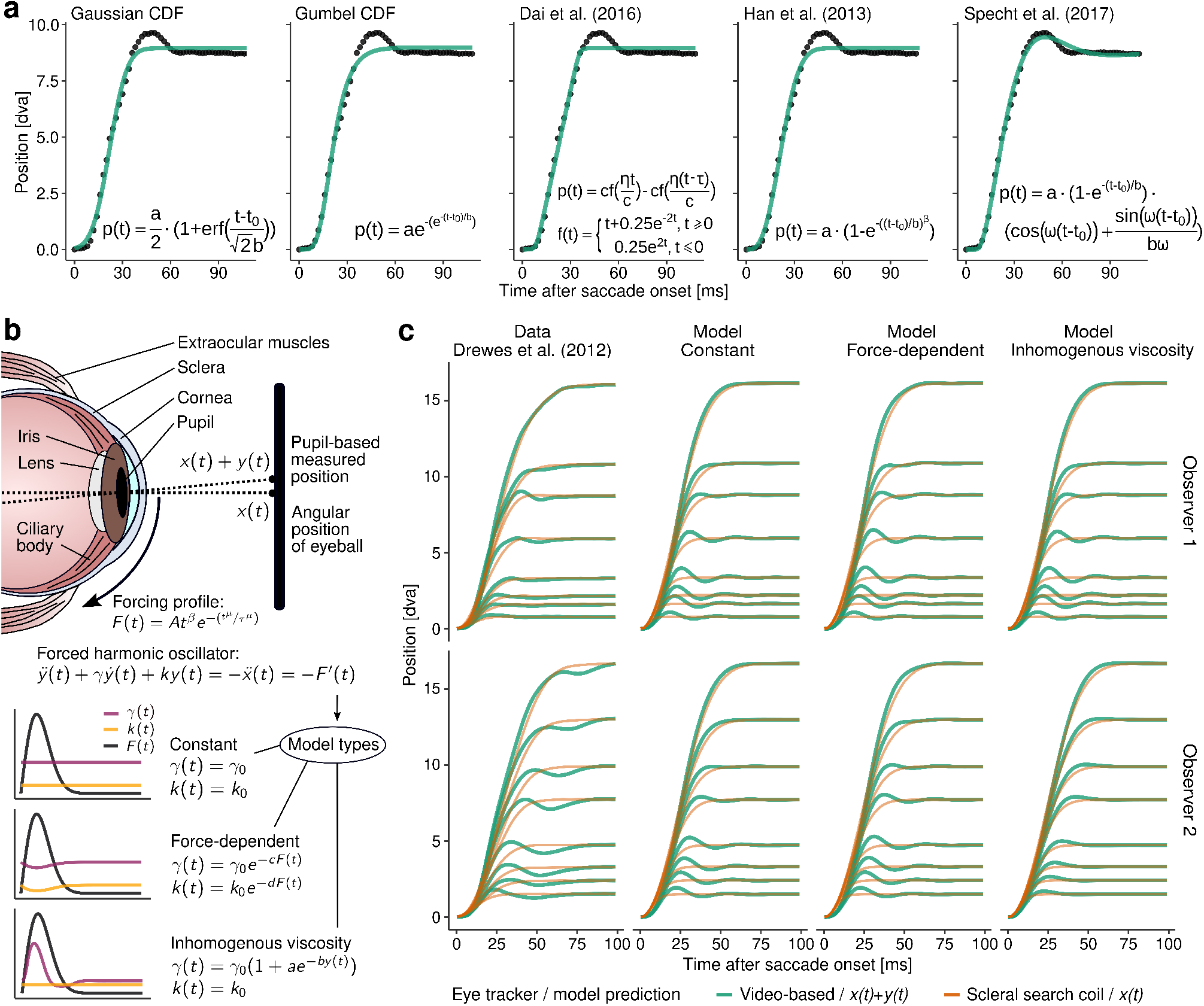
Modeling saccades and post-saccadic oscillations. **a** Collection of models previously used to fit saccade trajectories. Using the Levenberg-Marquardt algorithm, these models were fitted to a saccade featuring a prominent one-phase PSO measured with a SMI Hi-speed eye tracking system at a sampling rate of 500 Hz [22]. **b** Simple illustration of the eye’s anatomy and the central assumptions of the model proposed by Bouzat and colleagues [37, 38]. Three types of models were proposed, in which the viscosity and elasticity parameters *γ* and *k* are either constant or vary depending on either the force *F(t)* exerted on the eyeball or the relative position of the pupil *y(t)* relative to the position of the eyeball *x(t)*. **c** Left column: Average saccades of two human observers measured by Drewes and colleagues [39] using video-based eye tracking (green lines) and scleral search coils (orange lines) simultaneously. Other columns: Predictions of the three types of models for pupil motion *x(t)*+*y(t)* (green lines) and eyeball motion *x(t)* (orange lines), each assuming one set of parameters for each observer and across all amplitudes. These sets of parameters were estimated using a custom grid-search procedure that determined the smallest sum of squares for the entire set of saccades by brute force.

That the harmonic oscillator might indeed be a reasonable and physiologically plausible model for describing the inertial movement of the pupil embedded in the iris (or the lens in the case of DPIs) relative to the rotation of the eyeball has been proposed by Bouzat and colleagues [37]. As illustrated in **Figure 2b**, they suggested that what pupil-CR video-based eye trackers measure is essentially a combination of two interdependent movements over time, namely the angular position of the eyeball *x(t)*, which is driven by a time-dependent forcing profile *F(t)*, and the position of the elastic parts of the eye relative to the eyeball *y(t).* This relationship is captured by the harmonic oscillator equation 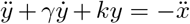 [37, Eq. 2], in which *γ* and *k* are the viscosity and elasticity constants, respectively. With respect to these parameters, the authors propose three different model variants: (1) *γ* and *k* may remain constant over time (constant model), (2) *γ* and *k* change as a function of exerted force (force-dependent model), or (3) *γ* changes as function of the iris’ deviation from its initial rest position while *k* remains constant (inhomogenous viscosity model). As noted in **Figure 2b**, two additional parameters quantify these relationships *– c* and *d* for force-dependent models or *a* and *b* for viscosity models, respectively. In all model variants, 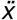 represents the force received by the iris (i.e., the acceleration of the eye-ball) which is equal to the derivative of *F(t)*, that is, 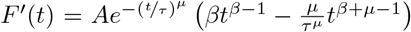 [38, Eq. 11], where parameter *A* relates to forcing strength and *β* and *μ* relate to the peak velocity and skewedness of the resulting velocity profiles. For convenience, the parameter τ can be substituted with 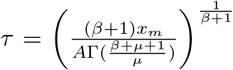 [38, Eq. 6], where Γ is the gamma function [45, Eq. 6.1.1] and *x*_*m*_ is the saccade amplitude. To allow the reader to assess the individual contributions of the model parameters, we implemented an interactive application which can be accessed at https://richardschweitzer.shinyapps.io/pso_fitting_example/. Finally, given all the above parameters and assuming an initial state of 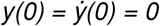 where the eye is at rest, the above differential equation for the harmonic oscillator can be solved numerically for a time sequence *t* (from 0 to an arbitrary value *t*_*max*_) to compute the relative movement of the pupil *y(t)*. To estimate the pupil’s entire trajectory, *y(t)* must be combined with the eyeball’s movement *x(t)*, which is defined as 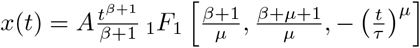 [38, Eq. 4], where _*1*_*F*_*1*_ is a confluent hypergeometric function [38, Eq. A1]. In order to facilitate the application of this procedure, we implemented it in the programming language R using the *deSolve* package for solving initial value problems [46]. It is publicly available at https://github.com/richardschweitzer/PostsaccadicOscillations and includes all model types and examples for fitting.

An advantage of the proposed model is that by just varying the saccade amplitude parameter *x*_*m*_, one set of parameters may produce entire families of saccades that have properties strikingly similar to empirical data: **Figure 2c** shows average saccades of two observers recorded simultaneously with video-based and search-coil measurements [39] along with the corresponding model predictions for pupil (*x*(*t*)+ *y*(*t*)) and eyeball (*x*(*t*)). Note that in this plot model predictions may not perfectly match the empirically measured saccades, which is owing to the fact that at this point only one set of parameters per model and observer was used to produce all saccade amplitudes (individual fits will be conducted in **section 2.2**). Yet, the model was able to reproduce some signature aspects of the empirical data. First, as first shown by Deubel and Bridgeman using a DPI [10], the movement of the eyeball precedes the movement of the pupil (or the lens) at saccade onset and also reaches its final position earlier. This delay, which occurs in all model variants, is the consequence of the inertial forces that act upon the elastic components of the eye. What follows, as we will later show, is that the saccadic peak velocity reached by the pupil should be larger than the peak velocity of the eyeball [6, 19].

This is also predicted by the model, because the velocity of the relative motion of the pupil, as it returns to and then overshoots its initial equilibrium position (i.e., *y* = 0), adds on to the velocity of the eyeball rotation. Second, both the data (clearly at least for observer 1; top row in **Figure 2c**) and the model predictions replicate the finding that the prominence of PSOs decreases with increasing saccade amplitudes [18, 20]. This could be explained by the idea that for shorter saccades it is more likely that extraocular forcing profiles directly overlap with the natural oscillation patterns of the iris, thereby producing a resonance-like phenomenon [37]. Similarly, a more tailed velocity profile, which is characteristic of larger saccades [34, 47], naturally contains longer deceleration periods and could thus allow the pupil to return to its equilibrium position in time for the end of the eyeball’s rotation. The model proposed by Bouzat, Del Punta and colleagues [37, 38] is thus capable of reproducing patterns of pupil and eyeball motion throughout the saccade, while resting on plausible physical principles. Yet, up to this point, it remains unclear how well this model actually corresponds to empirical data and what it could be used for by the eye-tracking community. Therefore, we will next critically investigate model predictions (**section 2.2**), especially the predictions of individual eyeball motion trajectories, and later provide an example how eye movement research could benefit from saccades that are authentically simulated by these kinds of models (**section 2.4**).

### 2.2 Predicting eyeball motion from pupil motion

One of the exciting possibilities that this model opens up is that, by fitting individual saccadic trajectories measured with video-based eye tracking, it may be possible to infer the unobservable motion of the eyeball from the observed motion of the pupil. In other words, by fitting the model to approximate *x(t)*+*y(t)*, we automatically extract the parameters for describing *x(t)*. The model has so far only been used to generate families of saccades [37, 38]. Here we will attempt the modeling of single saccades and evaluate the goodness of fit for pupil and eyeball predictions by comparisons with ground truth data.

Fitting the proposed models outlined in **section 2.1** to actual saccades requires the detection of those saccades first. We applied the widely used Engbert-Kliegl algorithm (see **section 2.3** for a detailed description) with a velocity threshold multiplier of 5 and a minimum duration of 15 samples, and only considered those saccade candidates that could be detected in both video-based and search-coil data. We fitted position profiles of saccades in a time window of 120 ms starting at *t* = 0, that is, the detected saccade onset in the search coil data, where the eye’s position was set to 0 dva. Considering the complexity of the model, fitting the model can prove a difficult task, as it may be difficult to find global minima or, in other words, the truly best fitting model parameters. More specifically, optimization algorithms will most likely find local minima which are heavily dependent on the supplied starting parameters [48]. To mitigate this problem, we used two procedures that worked reasonably well. First, one can specify an entire grid of starting parameters and run an optimization algorithm, such as the Levenberg-Marquart algorithm included in the R package *minpack.lm* [49], for every single starting parameter combination. For each run, the resulting estimates and the goodness of the fit are saved. After that, one may choose the result that best fits the data or has the most plausible parameter estimates. Second, one may also perform a two-step procedure. In the first step, the best starting parameters are determined by a grid-search (or random-search) approach, in which the best model parameters are determined by brute force in a rather coarse manner. An easy-to-use implementation is for instance provided by the *nls2* R package [50]. In the second step, these brute-forced model parameters can be refined when supplied as starting parameters to an optimization algorithm. An obvious disadvantage is that both of these approaches are computationally costly, especially as in each iteration (at least) one ordinary differential equation has to be solved numerically. Yet, these iterations can often be easily parallelized [e.g., 51], and parameters have limited ranges of plausibility (i.e., *β*: 1–1.3, *μ*: 1–4, *A*: 0.02–0.08, *γ*: 0.05-0.20, *k*: 0.01-0.10; see [38] for further details).

**Figure 3a** shows the best parameter estimates across saccade amplitudes for both observers and three model types, clearly suggesting that only one set of parameters is unlikely to account for an observer’s entire saccadic main sequence. For instance, larger saccade amplitudes are better fitted by higher values of *A*, indicating an increase of mechanical forcing strength. Moreover, larger amplitudes are associated with lower values of *γ*, suggesting that viscosity decreases, thus leading to more pronounced PSOs. On the one hand, it may well be that viscosity and elasticity change depending the applied external force, as elastic parts of the eye may be not only rotated, but also deformed by that force [37]. On the other hand, as the model has been shown to underestimate PSO amplitudes at larger saccade amplitudes (**Figure 2c**), lower values of *γ* may simply be needed to match the larger-than-predicted PSO amplitudes present in the data. Fitting individual saccade trajectories (compared to using one set of parameters for an entire family of saccades) thus improves the model representation of PSOs that occur at larger saccade amplitudes: Modeled trajectories for 16-dva saccades shown in **Figure 3c** are in fact strikingly similar to those found in previous studies [e.g., 20, Figs. 8 and 10]. It is also quite evident that high values of *μ* are especially prominent at short saccades, resulting in brief, high-velocity forcing profiles and leading (as we will show in **Figure 3d-e**) to an underestimation of the eyeball’s actual rotation duration at very short saccade amplitudes.

**Fig. 3:**
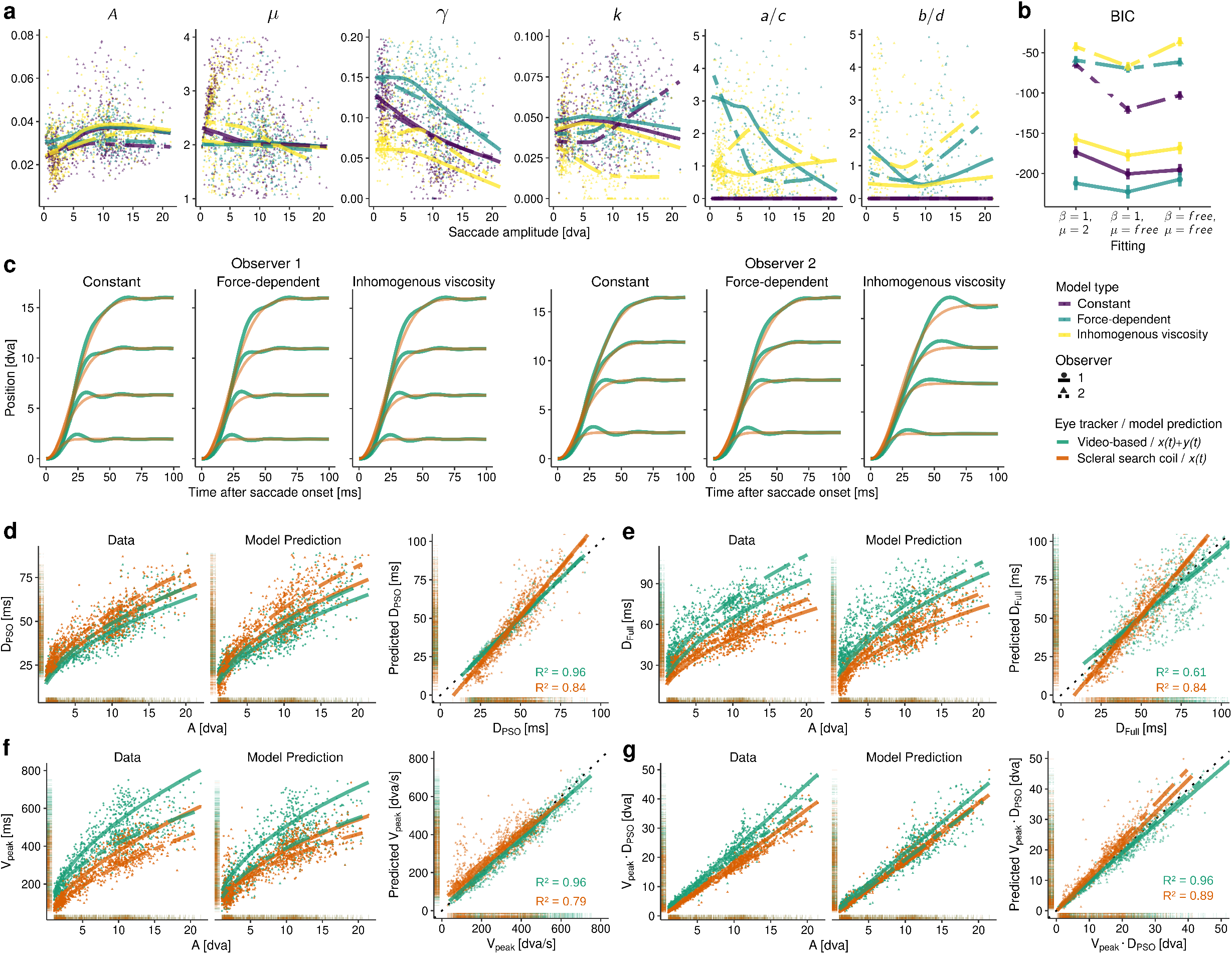
Fitting individual saccade trajectories to predict eyeball motion from pupil motion. **a** Parameter estimates for best-fitting models of individual saccade trajectories (extracted from data by Drewes and colleagues [39]) for observers 1 (circles, solid lines) and 2 (triangles, dashed lines). Colors indicate the three model variants: Constant (purple), Force-dependent (green), and Inhomogenous viscosity (yellow). **b** Goodness of fit, as measured by Bayes information criterion (BIC), for the three model types and three options for treating parameters *β* and *μ*: Setting both to their proposed default values [37], adding *μ* as a free parameter, or adding both *β* and *μ* as free parameters. **c** Results of individual fits, using the mean best fitting parameters for each observer, model, and saccade amplitude. Predictions are shown for pupil motion (green lines) and eyeball motion (orange lines). **d-g** Main sequence relationships of original and modeled saccades for observers 1 and 2. Model predictions for each saccade were produced by the best model, as indicated by the lowest BIC. Color indicates the measurement type: Video-based tracking and pupil motion (green) vs. scleral search coil tracking and eyeball motion (orange). Right figures in each panel show the statistical relationship between original saccade metrics and those predicted by the model.

As parameters *A, β,* and *μ* all manipulate the shape of the forcing profile in a rather similar manner [38, Fig. 2], is it necessary to include all of them as free parameters? Measures of model parsimony (**Figure 3b**) suggest that it is reasonable to include *μ* as a free parameter – especially if the Constant model is used – but not necessarily *β*. This parameter can be safely set to *β* = 1, especially as values of *β* < 1 would lead to infinite acceleration and are therefore meaningless [38]. With respect to the question which model type most parsimoniously fits the data, our analysis suggests to choose either the constant or the force-dependent model. If it is crucial to minimize computing time, then it is advisable to choose the constant model, as its lower number of free parameters reduces run times of the fitting procedure by at least a factor of 3.

To what extent is the model capable of reproducing an observer’s saccadic main sequence? To answer this question, we let each saccade’s best-fitting model predict a position profile, and performed velocity-based saccade detection on both the predicted and the real saccades to extract metrics like *PSO*-defined and *Full*-defined saccade duration (*D*_*PSO*_ and *D*_*Full*_, see **Figure 1b**), peak velocity (*V*_*peak*_) and the product of the latter (*V*_*peak*_ · *D*_*PSO*_). Note that most saccade detection algorithms use two-dimensional position data (i.e., [*x*,*y*] over time), while the model discussed here uses one-dimensional position data (i.e., position change of time). As long as the direction vector of the original saccade (relative to the saccade starting point [*x*_*on*_, *y*_*on*_]) is known, model predictions can always be transformed back to two-dimensional data (**Figure 1a**), thereby even reproducing the original saccade’s curvature. When looking at the capability of the model to accurately reproduce saccade metrics, two aspects of the fit may be evaluated, namely (1) how well the model is able to fit pupil-based signals of saccades and post-saccadic oscillations, and (2) how well the model can infer the latent metrics of eyeball rotation, which are otherwise unobservable with video-based measures. Regarding the first aspect, the results show that the model is capable of matching both *PSO*-defined saccade durations (**Figure 3d**) and peak velocities (**Figure 3f**) with high coefficients of determination. The exact shape of the PSO, however, could not be represented with such precision, as indicated by the on average veridical, but considerably weaker relationship between actual and predicted *Full*-defined saccade duration (**Figure 3e**). With respect to the second aspect, results suggested that the model inferred duration and peak velocity of eyeball motion reasonably well, even though systematic prediction errors became apparent. Saccade duration was underestimated at very small amplitudes and overestimated at larger amplitudes (**Figure 3d**), whereas saccadic peak velocity was overestimated at very small amplitudes (**Figure 3f**). These prediction errors could be explained by the assumption that the underlying forcing profiles of coil-measured short saccades may actually be quite different from the profiles assumed in the model, as short saccades or microsaccades have in fact been shown to exhibit dynamic overshoots, as well [16, 18, 19]. As the current model does not incorporate this aspect, forcing profiles could become increasingly brief and rapid at small amplitudes (as indicated by increased estimates of

*μ* in **Figure 3a**) to accurately fit the prominent PSOs in the video-based pupil data. Note that, due to their small size and low velocity, it can be difficult to detect coil-measured PSOs using velocity thresholds common for video-based systems [e.g., 10–30 dva/s, 52]: In the present analysis PSOs were only detected in less than 3% of all coil-measured saccades (as compared to 90% of all video-measured saccades), which is why we did not consider separate analyses for different saccade definitions in search-coil data. As the data has been made publicly available for more in-depth analyses (https://osf.io/n36fx/), it may be an interesting avenue for future research to refine the assumptions of the proposed model [37, 38] by applying more physiological plausible forcing profiles based on models of motoneural activity [4, 12].

Finally, given that the fitting procedure described above is quite complex, one could ask whether there may be simple statistical heuristics that a researcher can use to approximate the metrics of eyeball motion from the metrics observed in video-based eye tracking. As already suggested by Deubel and Bridgeman [10], there is a very tight relationship between the time of the first peak of a PSO and the offset of a coil-measured saccade, as indicated by high values of *R*^*2*^ and regression slopes close to *β* = 1 (**Figure 4a**). More specifically, significantly positive intercepts in the two observers shown here indicate that *PSO*-defined saccades may be 1–6 ms shorter than coil-measured saccades, suggesting that the onset of the PSO is a valid criterion for the offset of eyeball rotation. In contrast, the offset of the PSO has little agreement with the offset of coil-measured saccades as indicated by large systematic overestimation of saccade duration, especially at longer durations, and considerably larger variance (**Figure 4b**). As already evidenced by previous studies [6, 18, 19], a similar – but in this case quite reliable – overestimation occurs for saccadic peak velocity (**Figure 4c**) and (to a lesser degree) for saccadic mean velocity (**Figure 4d**). Note that fitting video-based saccade data with the described model above can also produce accurate velocity estimates for the underlying eyeball motion (**Figure 3f**).

**Fig. 4:**
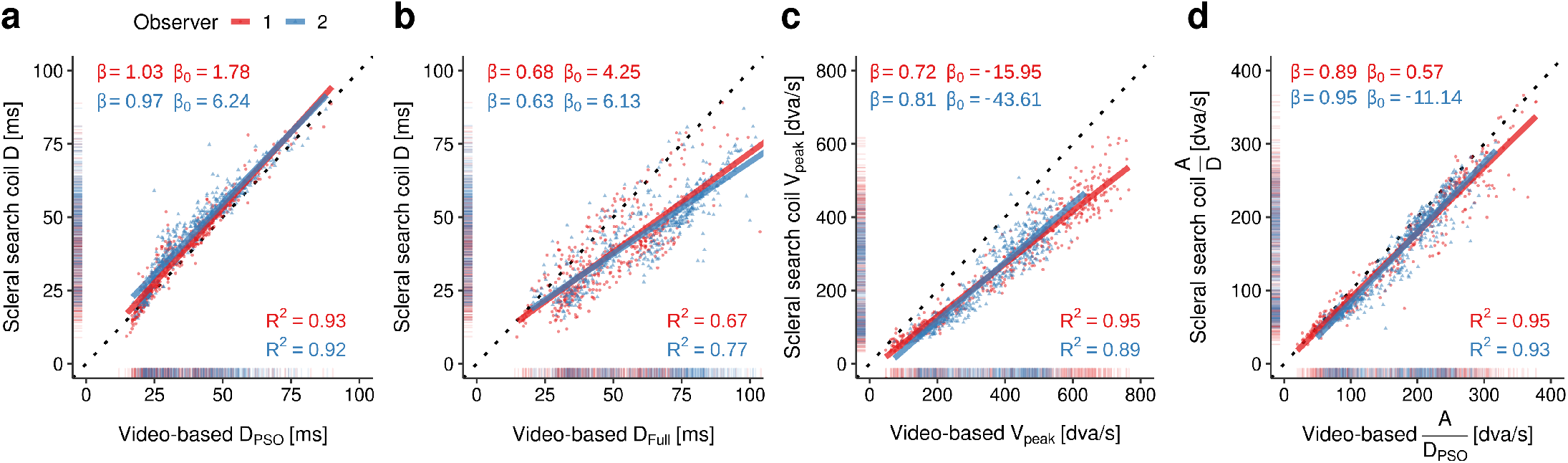
Statistical relationships between saccade metrics (**a**: *PSO*-defined duration, **b**: *Full*-defined duration, **c**: Peak velocity, **d**: mean velocity defined as the quotient of amplitude and duration) measured by video-based and scleral search-coil systems, based on data from two human observers collected by Drewes and colleagues [39]. Linear regressions (solid lines) were performed separately for observers 1 (red) and 2 (blue) on a total of 960 saccades of varying directions. *β* and *β*_0_ indicate slope and intercept, and *R*^2^ the coefficient of determination of each fit.

These results indicate lawful relationships between pupil and eyeball motion, suggesting that optical measures of gaze position, previously considered “not suitable for the analysis of saccade dynamics (main sequence)” [10, p. 536], are in principle able to provide interpretable estimates of ‘true’ saccade metrics. They also suggest that it may really be worthwhile to detect PSOs, as *PSO*-defined saccades offsets have proven to be not only more accurate, but also more precise measures for saccade duration. The next two chapters will thus be dedicated to algorithms that allow eye tracking researchers to detect PSOs as part of of saccades.

### 2.3 Direction-based and velocity-based detection of saccades and post-saccadic oscillations

There is extensive literature on automated saccade detection. Of the vast number of algorithms available, each has specific strengths and capabilities, as well as varying degrees of complexity and accuracy. Whereas earlier detection algorithms relied on absolute or relative thresholds for position, velocity, or acceleration [e.g., 40, 53, 54], more recent algorithms involve covariance [55], binocular correlation [24], particle filters [56], Bayesian inference [57], or Markov models, Kalman filters, and spanning trees [58, 59], to name just a few. Andersson and colleagues [21] have evaluated the performance of ten different algorithms, among which two algorithms were specifically dedicated to the detection of PSOs [5, 22]. These two algorithms exhibit high detection performance, one of them even in the face of smooth pursuit [22], yet they are relatively complex. Here, we instead describe an comparably simple, sequential approach, in which we first detect saccade candidates using the popular Engbert-Kliegl (EK) algorithm for microsaccade detection [40, 41], then identify clusters of such candidates, and fianlly detect the PSO based on the direction of the first detected saccade (**Figure 5a**).

**Fig. 5:**
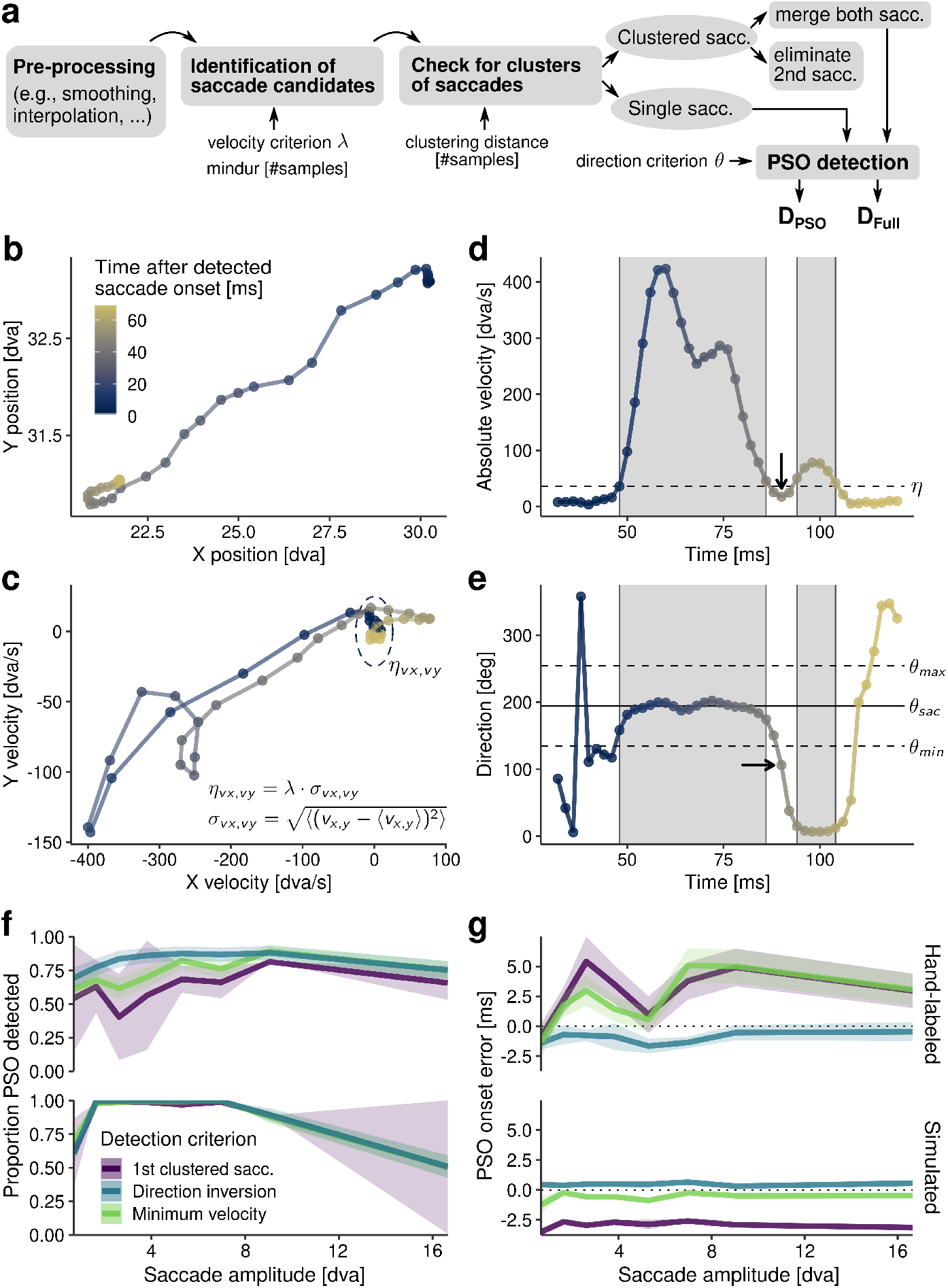
Velocity- and direction-based detection of saccades and post-saccadic oscillations. **a** The proposed pipeline for PSO detection. *D*_*PSO*_ and *D*_*Full*_ refer to the saccade definitions in Figure 1b. **b** Gaze position data of a saccade taken from a hand-labeled dataset recorded using a SMI video-based eye tracking system [also used in 21–23]. **c** Two-dimensional velocity space of the same saccade. The dashed ellipse indicates the velocity thresholds η_*vx,vy*_ computed based on the median-based standard deviations σ_*vx,vy*_. **d** Absolute sample-to-sample velocity in degrees of visual angle replotted as a function of time. Gray-shaded areas represent saccade intervals in which samples are above the velocity threshold η (dashed horizontal line). The arrow indicates the minimum velocity of the merged interval. **e** Sample-to-sample direction in degrees relative to the overall saccade direction *θ*_*sac*_ (solid horizontal line) and defined direction thresholds *θ*_*min*_ and *θ*_*max*_ (dashed lines). The arrow indicates the first sample to cross the direction threshold (±60 degrees), which coincides with the minimum velocity of the merged interval for this particular saccade (arrow in panel c). **f** Proportion of PSOs correctly detected in a hand-labeled (upper row) and simulated (lower row) datasets. Color indicate the three proposed detection criteria: Offset of the first saccade in a cluster of two saccades (purple), first direction sample to deviate from the overall saccade direction (blue), and sample with the lowest velocity of the entire saccade cluster (green). **g** PSO onset detection error, defined as the time between the coded PSO onset (upper row: human hand-labellers, lower row: model-based simulation) and detected PSO onset. Positive values indicate that detected onsets occur later than coded onsets. Shaded areas indicate ±*SEM*.

The EK algorithm is a velocity-based detection method. Gaze position data is transformed into a two-dimensional velocity space, usually using a five-point moving average to reduce sample-to-sample noise. Even though only data of a single saccade is shown in **Figure 5b**, entire sequences of saccades can be submitted to the procedure. Based on the median-based standard deviation σ_*vx,vy*_ of all available samples and a user-defined threshold multiplier λ (usually taking values of 5–10), an elliptic velocity threshold η_*vx,vy*_ is computed (**Figure 5c**). A velocity sample above this threshold is classified as belonging to a saccade, according to the test function 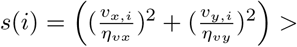 [60, Eq. 4]. As velocity may cross the threshold simply due to noise, sequences of above-threshold samples must have minimum length (*mindur*, defined as number of samples) to be accepted as saccade candidates. As shown in **Figure 5d**, this approach based solely on velocity may result in the detection of two, clustered saccades, instead of detecting one saccade with a PSO. In fact, large PSOs may well reach peak velocities of 100 dva/s and durations of 40 ms [5], thus easily satisfying standard criteria for (micro-)saccade detection.

How should the researcher deal with this situation? In **Figure 5a**, we propose a simple processing pipeline: After optional pre-processing of the data, which may contain filtering, interpolation (note that in-built velocity smoothing in the EK algorithm only works for uniformely sampled data), and removal of blink artifacts [e.g., 61], saccade candidates are identified. Even though we chose the EK algorithm (due to its prevalence in the field), any saccade detection algorithm may be used at this stage. To determine whether clusters of saccades (as in **Figure 5d**) are present in the data, we check for two or more saccade candidates with only a small number of below-threshold samples in between. This clustering distance can be chosen by the user. Although fixation durations below 80 ms are quite rare, they may occur [62], which is why we recommend to minimize the clustering distance (e.g., 5–20 samples, depending on the sampling rate) to avoid the erroneous clustering of separate saccades occurring in close succession. If clustered saccades are detected, there are essentially two options. First, one may discard the second part of the clustered saccade (i.e., the PSO), leading to a saccade offset that is defined as the time the velocity first falls below the defined threshold and may thus be closely related to the peak of the PSO. Second, one may choose to merge the two clustered saccade candidates and subsequently perform a dedicated PSO detection. For PSO detection, we propose two possible criteria: The minimum-velocity criterion determines the lowest sample-to-sample velocity in the entire merged saccade interval (not including the first and last velocity samples; **Figure 5d**), and the direction-inversion criterion detects the point where the sample-to-sample direction first significantly deviates from the overall saccade direction *θ*_*sac*_, that is, when crossing a direction threshold defined by *θ*_*min*_ and *θ*_*max*_ (**Figure 5e**). This direction threshold can be adjusted by the user by setting the parameter *θ* (*θ*_*min,max*_ = *θ*_*sac*_ ± *θ*). It becomes clear in **Figure 5b** [but see also 63, Fig. 20] that a consequence of PSOs is the brief inversion of direction relative to the overall saccade direction, as defined by the mean direction of all saccade samples from the respective saccade onset position (**Figure 1a**). Code for this procedure (and the one described in **section 2.4**) can be found at https://github.com/richardschweitzer/PostsaccadicOscillations.

How does the proposed detection pipeline perform, and are there differences between the three detection criteria? To answer these questions, we performed a validation procedure using 300 hand-labeled saccades with PSOs [21–23]. As manual coding may be prone to subjective biases [64], however, we propose a second validation approach: Based on the introduced modeling approach (**section 2.1**), we simulated saccades with PSOs along with fixation data (see next **section 2.4**) with objective labels for fixations, saccades, and PSOs, thus providing a model-based ground truth. Results suggest that PSOs were correctly detected at high rates in both hand-labeled and simulated saccades; the best detection criterion turned out to be the inversion of direction (**Figure 5f**). Clearly, simply relying on the detection of a saccade cluster (in order to subsequently remove its second saccade) proved a suboptimal strategy, as saccades and PSOs are not necessarily detected as two separate events. What is more, PSOs may not have sufficient velocity and duration to be detected as an event to begin with. Furthermore, simply detecting minimum velocity within a saccade cluster had disadvantages, as sample-to-sample velocity signals can be extremely noisy, especially when measured with high sampling rates [43]. We also found a clear tendency in both datasets that detection performance decreased at very small and very large saccade amplitudes. This is most likely owing to the fact that very short saccades and microsaccades, as well as their PSOs, produce only weak velocity signals which may well remain undetected within normal tracking noise. Large saccades, in turn, produce smaller PSO amplitudes that may be missed due to their reduced prominence [20]. Direction inversion also proved to be the most accurate criterion for PSO onset, both for hand-labeled and for simulated saccades, exhibiting absolute errors of only 1 ms across saccade amplitudes (**Figure 5g**). In contrast, results for the two velocity-based criteria look less convincing. On the one hand, they show delayed PSO-onset detection by up to 5 ms relative to human coders. On the other hand, with simulated data, velocity-based criteria detect PSO onsets prior to their actual onset, which is consistent with theoretical assumptions, as velocity should fall below the velocity threshold defined by the EK algorithm before it approaches zero and before the direction inversion takes place. We can speculate that human coders might have used different subjective criteria; in fact, it seemed that they would often define PSO onset as occurring slightly before the first peak of the PSO (cf. **Figure 6d**). Regardless of this mismatch, detection of PSOs, based on saccade clustering and the application of minimum-velocity or (even better) direction-inversion criteria, is feasible and may well be used as an extension to various saccade detection algorithms.

**Fig. 6:**
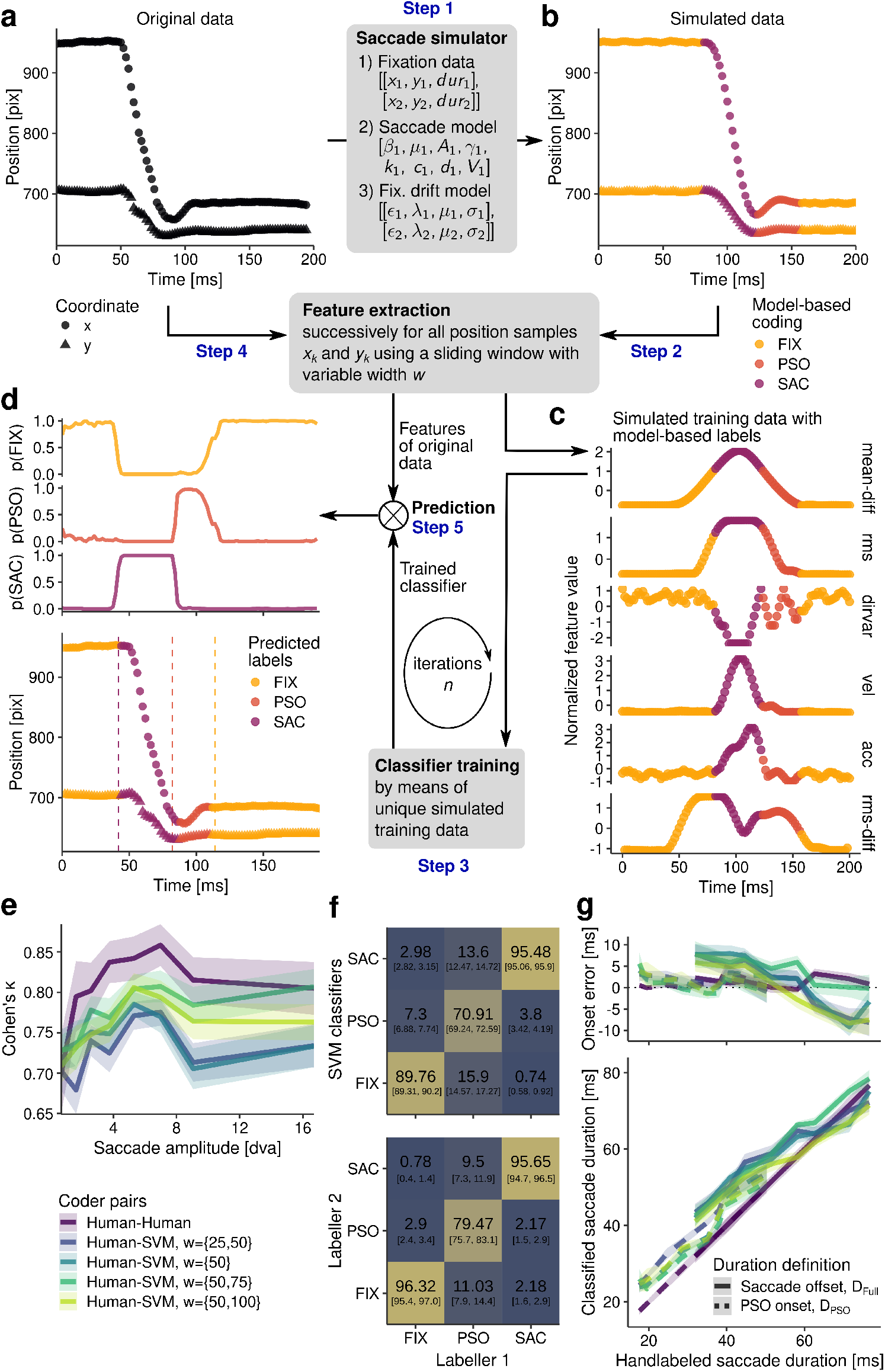
Accurate detection of saccades and PSOs using linear classifiers trained on simulated data. **a** Original, unlabeled saccade data. **b** Using a saccade simulation algorithm, incorporating models for saccades and PSOs [37, 38], as well as models for fixational drift [65], training data with class labels is produced. **c** Feature extraction is performed for every single data sample of the simulated data by integrating over a window with a width of *w* [66]. These simulated features are used to train a linear support vector machine [67]. **d** The same feature extraction is performed on the original unlabeled data. Based on the resulting features, the previously trained classifier may predict class labels (dashed lines indicate hand-labeled event onsets). **e** Cohen’s *κ* as a metric for inter-rater agreement as a function of saccade amplitude. **f** Confusion matrices (upper: human vs. classifier, lower: human vs. human), computed across all saccades and window sizes. **g** Top row: PSO-onset and saccade-offset detection errors (classified onset time – handlabeled onset time) produced by classifiers and human experts. Bottom row: Relationship between hand-coded and classifier-coded saccade durations for duration definitions *D*_*Full*_ and *D*_*PSO*_ (Figure 1b).

### 2.4 Simulation of saccades and post-saccadic oscillations for classification-based detection

To date, the best saccade detection algorithms use machine learning techniques [66, 68] or even deep neural networks [69, 70], thereby achieving human-like classification performance. These algorithms have to be trained on annotated eye movement data, usually provided by hand-labelling performed by human experts. Although this procedure is widely considered the gold standard [64], it is extremely time-consuming, especially when high inter-rater reliability has to be reached despite subjective biases. This issue could be alleviated if saccade and fixation data could be generated by models in a realistic manner. We take this approach here as it has evident advantages: Data produced by a model has an actual, mathematically verifiable ground truth, and the modeler retains full control over the signal’s quality, so that, for instance, effects of noise or missing samples can be systematically investigated [32, 42].

To allow for such investigations, we implemented a multi-purpose saccade simulator (**Figure 6a-b**). Sequences of saccades can be reproduced if only approximate sequential fixation positions (and optionally fixation durations) are known. For each saccade, eight parameters have to be supplied to describe its trajectory (see **section 2.1** for a description of these model parameters, while *V* is a boolean value indicating whether the viscosity-based variant of the model should be used). Note that these parameters could be in principle the same for the entire sequence of saccades. For each fixation, four parameters are specified to generate fixational drift patterns that are common to fixation periods [28, 71]. Here we used a self-avoiding random walk model by Engbert and colleagues [65] to approximate these drift patterns. This model assumes a two-dimensional quadratic lattice with a width of *L* pixels (that should amount to approximately 2 dva in a given setup) in which each field has a certain activation *h*. In our version of the model, each field is preactivated with a random number drawn from a Gaussian distribution with mean *μ* and standard deviation *σ* (where negative values are set to 0). In addition to varying activations, the lattice is goverened by a quadratic potential with the formula 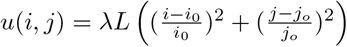 [65, Eq. 3], where *i*_*0*_ and *j*_*0*_ represent the lattice’s center coordinates and *λ* is the steepness of the potential. Starting at (*i*_0_, *j*_0_), a random walker moves around in the lattice, always choosing its future position based on the lowest current activation and potential: (*i*′, *j*′) = *argmin* {*h*_*kl*_ + *u*(*k, l*)}, where *k* = *i* ± 1, *l* = *j* ± 1 [65, Eq. 4]. Crucially, each visited position *h*_*ij*_ is increased by 1 in its activation, so that the walker will only return to this position after some time, that is, after the activation has decayed with *h*_*kl*_ → (1 – *ε*) · *h*_*kl*_ [65, Eq. 2] per step (each step being one data sample). To model drift movements on a miniture scale during a fixation interval, the user simple has to specify the decay rate *ε*, the steepness of the gradient *λ*, and the distribution (*μ*, *σ*) for the pre-activation of the lattice grid. As these drift movements can (depending on the specified sampling rate) nevertheless result in comparably high velocities, an optional five-point moving average can be used to reduce noise (**Figure 7**).

**Fig. 7:**
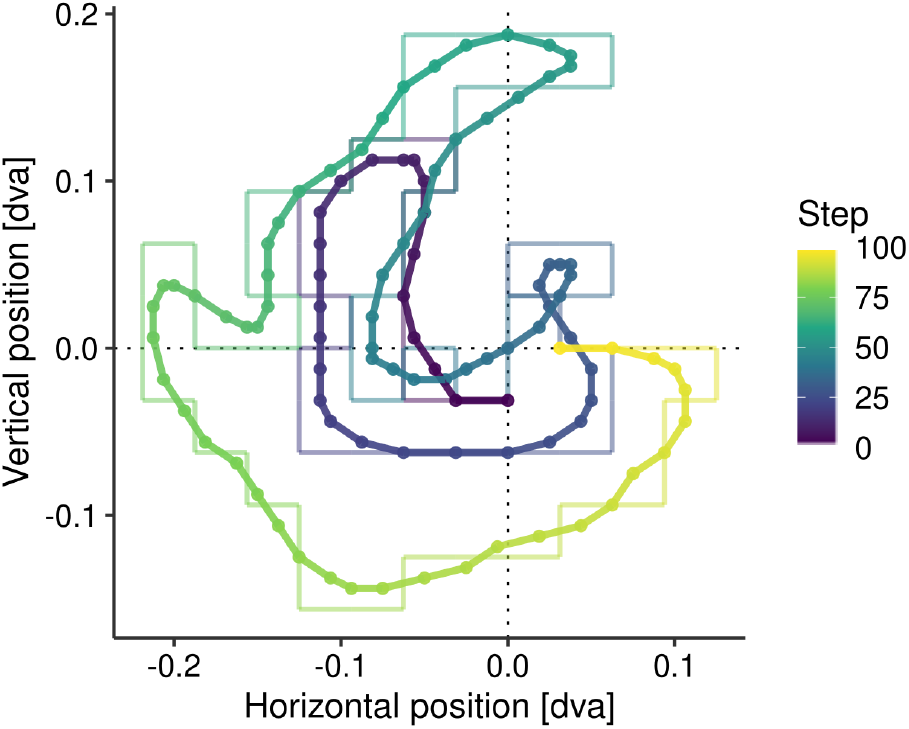
Fixational drift patterns produced by the self-avoiding walk model proposed by Engbert and colleagues [65]. To reduce the discreteness of the steps (transparent lines) and reduce the signal’s velocity, we applied a five-point moving-average smoother. To approximate physiological ocular drift even better, the number of steps per time interval and steps per degree of visual angle may be adjusted, as well as the model paramaters decay rate *ε*, gradient steepness *λ*, and mean and standard deviation of the grid initialization distribution (*μ*, *σ*).

By simulating fixational drift and saccades, the user can produce realistically looking eye tracking data, which can then be used to evaluate detection algorithms (**Figure 5f-g**) or train models for classification. Here we propose a novel detection procedure based on simulation and classification that involves five major steps and is illustrated in **Figure 6**. As a first step, we use the saccade simulator described above to generate saccade and fixation data that is similar, but not necessarily identical to the original data (**Figure 6a-b**). Importantly, this simulated data includes model-generated event labels (*FIX*, *PSO*, *SAC*) that will later serve as the dependent variable in the model training phase. As a second step, the simulated gaze position signal (*x*, *y*) at each time point *k* (**Figure 6b**) must be transformed into a vector of meaningful features that can be used as independent variables. To achieve that, we followed an approach proposed by Zemblys and colleagues [66] that integrates position data using a temporal window with a width of *w* samples. Note that *w* may vary depending on the type of feature. For instance, the root mean square (*rms*) feature at time point *k* was computed based on all samples in the range 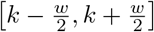, whereas the difference of mean position (*mean-diff*) was defined as the absolute difference between the mean gaze position in [*k* − *w, k*] and the mean gaze position in [*k, k* + *w*]. The features absolute velocity (*vel*) and acceleration (*acc*) used a 12-ms window, in order to be able to represent brief saccades [5], whereas the variance of sample-to-sample directions (*dirvar*), that is, 1 – the mean resultant length of the circular distribution [72], used a window of 22 ms to adequately capture direction changes. After this step, we extracted 13 features per window size from simulated data [see 66, for a full list of features], covering a wide range of possible indicators of fixations, saccades, and PSOs (**Figure 6c**). Now that both event labels and features (each ideally centered and scaled) are available, they can be used to train classifiers. The third step is thus the model training phase. Here, we used a linear support-vector machine (L2-regularized, L2-loss, primal), as implemented in the *liblineaR* package [67], but note that a wide range of other (potentially more effective) classifiers have been successfully employed for this task [68]. In the fourth step, the same feature extraction procedure, which has previously been used on simulated data, is now applied to the original, unlabeled data. Finally, in the prediction step, the trained classifier can be used to determine an event label for each sample of the original data. Note that this procedure can be repeated a number of times, each time with slightly different training sets, to make individual predictions more robust or to determine samples with ambiguous class attributions (**Figure 6d**, upper rows).

One may now ask how well this procedure performs compared to manual coding performed by human experts. To answer this question, we ran the proposed procedure on 300 hand-labeled saccades with PSOs [extracted from a dataset used by 21–23], each time choosing the most likely of 100 labels predicted by uniquely trained classifiers. To allow for a fair comparison between human coders and classifiers, we introduced the agreement between the two human coders, that had previously manually labeled the data, as a reference. As indicated by Cohen’s *κ* (i.e.,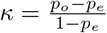, where *p*_*o*_ is the observed proportionate agreement and *p*_*e*_ the expected proportionate agreement, if labellers rated based on chance [73]), these two coders do not agree completely – most likley due to individual criteria – but show *κ* around or above 0.8, which is generally considered “almost perfect” [74, p. 165], although their agreement slightly decreased at very short and large saccade amplitudes (**Figure 6e**). Importantly, classifiers were able to reach near-human coding performance, even though this depended on the windows sizes that were used. By trend, using short, or only one window size led to lower performance, whereas maximum classification accuracy was found with window widths ranging from 50 to 100 ms. Note that each sample was classified independently from its surrounding samples. The introduction of transition probabilities, minimum durations for events, or even simple rules, such as “PSOs do not occur in the midst of a fixation period” could thus further increase coding performance. In addition, it was shown that, compared to linear support-vector machines, decision trees and random forests were even more suitable classifiers for this kind of task, reaching up to *κ*=0.9 when being trained and evaluated on hand-labeled data [68]. It is thus well possible that the procedure explained here could be further optimized. **Figure 6f** shows the confusion matrices, which indicate in which cases humans and classifiers (upper table), as well as humans and humans (lower table) disagreed: Clearly, whereas saccades and fixations were extremely well separated, most mismatches occurred when coding PSOs. This pattern was present for both human-classifier and human-human rating agreements, suggesting that it is inherently difficult, even for human eye tracking experts, to determine PSO onset after saccades, as well as fixation onsets after PSOs. In addition, it is quite likely that detection criteria differed not only between human labellers, but also between the hand-labeling procedure [for the custom Matlab-based GUI, see 21, Fig. 1] and our model-based simulations. As suggested by **Figure 6d**, manually coded PSO onsets may not necessarily coincide with the first peaks of PSOs which would be determined by the saccade simulator. Similarly, PSO durations may be overestimated by algorithms, as these may still pick up spurious velocity, acceleration, or direction signals that a human coder would disregard. To evaluate these inherent differences, we compared saccade metrics resulting from classification with those resulting from manual coding. Indeed, saccade duration was slightly overestimated by virtually all classifiers (**Figure 6g**). This was caused by an earlier detection of saccade onsets (by on average 2 ms), as well as a later detection of PSO offsets (*D*_*Full*_ definitions) by up to 8 ms, most likely due the classification routine’s more fine-grained resolution. In fact, an identical result was found by Andersson and colleagues [21], who showed that two previously proposed PSO-sensitive saccade detection algorithms [5, 22] overestimated PSO duration (compared to human coders) to a similar extent. Yet again, PSO onset (*D*_*PSO*_ definitions) proved a more reliable criterion for saccade offset, as estimates of saccade durations, as well as onset errors were smaller and less variable (**Figure 6g**).

Here we have shown that it is not only well possible, but also highly efficient to use simulated saccades, PSOs, and fixations as training data for classification-based approaches of automatic detection of events in eye tracking data. All of these relatively recent classification-based approaches [66, 68–70] assumed manually coded eye tracking data as a ground truth, even though the validity of this type of reference has been questioned, as empirical data can indeed be ambiguous and even human experts may have individual detection criteria [64]. Thanks to sophisticated biophysical models [37, 38, 44, 65], simulation of eye movement data may be a feasible alternative – not only because of mathematical tractability, but also because of greatly reduced effort for the researcher.

## 3 Conclusions

In the first part of the chapter, we have introduced physiologically plausible models [37, 38] capable of describing relative pupil and eyeball motion (**section 2.1**) that – simply based on biophysical premises – provide an account for post-saccadic oscillations (PSOs). Despite their high relevance to the field and fitting well with previous empirical results, these recently proposed models have not yet had any impact on the eye tracking community (at least not to our current knowledge). One could speculate that this is owing to the fact that the models in question have neither been implemented in a ready-to-use manner, nor have their predictions been evaluated in a critical comparison with suitable empirical data. To bridge this gap, we provided openly accessible code which can be used to generate and fit saccade trajectories with PSOs. Capitalizing on co-registered scleral-coil and video-based eye tracking data [39], we quantified to what extent models fitted on the pupil signal of a saccade could predict the eyeball motion of that very same saccade. We found that the model predicted eyeball motion with reasonably high accuracy, closely reproducing main sequence relationships of *PSO*-defined saccade duration and saccadic peak velocity for medium-sized saccades, whereas failing to do so for very short saccades. We speculate that this due to the forcing profiles of short saccades, which may also exhibit dynamic overshoots [14, 16, 18, 19]. It would be an intriguing avenue for future research to expand and improve the current models to more accurately match empirical saccade data. In fact, a near-perfect matching would mean considerable progress for the field, as that would allow for separate saccade definitions based on video-based data alone: On the one hand, definitions based on the motor commands executed by the extraocular muscles and, on the other hand, definitions based on the visual consequences induced by the movement of the elastic parts of the eye [11, 30]. At some point, these definitions may even be operationalized in future investigations of retinal and extra-retinal contributions to visual perception around and during saccades.

In the second part of the chapter, we described (and provided code for) algorithms that allow for the detection of PSOs, which may be helpful or even crucial for a number of applications. For one, we have shown that *PSO*-defined saccade durations closely match saccade durations measured by scleral search coils [cf. 10], follow the main sequence more tightly, and are less variable. Therefore, when close or conservative approximations of saccade offset are needed, detecting PSOs may drastically improve data quality. Examples of such applications include determining secondary saccade latencies [75–78], making sure that a presentation sequence has occurred strictly during a saccade [32], or measuring electrophysiological potentials, such as lambda waves, that are locked on saccade offset [79, 80]. While PSO detection based on direction inversion proved a simple, yet efficient solution, that can in principle be added onto most detection algorithms, saccade detection based on machine learning is the current state of the art. Using simulated saccade data with model-generated labels, practitioners in the field may be able not only to evaluate their custom detection methods, but also to train a wide range of statistical learning models without relying on manually coded data. While manual coding will remain the gold standard for various reasons, tests on such simulation data – due to their objectivity and tractability – may serve as proofs of concept and provide insights into how humans detect events in eye tracking data.

Finally, PSOs are prominent features of virtually any video-based eye tracking dataset. As they can be regarded as separate events, eye tracking researchers should make an informed choice as to whether they want to count them as being part of either saccades or fixations, and explicitely specify their approach. Here we raise awareness that the treatment of PSOs can make a real difference, not only conceptually, but also with respect to the interpretation of saccade metrics and experimental results. With this chapter and the methods specified in it, we hope to enable eye tracking researchers to make the most of their data by actively defining, modeling, and detecting saccades – not despite, but with the help of post-saccadic oscillations.

## 4 Acknowledgements

We thank Jan Drewes, Guillaume Masson, and Anna Montagnini for sharing their co-registered search-coil and video-based eye tracking data [39]. Furthermore, we thank Markus Nyström and Richard Andersson [21–23] and Anna-Katharina Hauperich and colleagues [24] for making their annotated datasets publicly available, as well as Ralf Engbert, Petra Sinn, Konstantin Mergenthaler, and Hans Trukenbrod for their R implementation of the Microsaccade Toolbox (http://read.psych.uni-potsdan.de/attachnents/article/140/MS_Toolbox_R.zip). We thank Antonio Sánchez Garcia for his help with the Phantom high speed camera and Sven Ohl and Wiebke Nörenberg for volunteering to provide feedback on an earlier version of the chapter.

R.S. and M.R. were funded by the Deutsche Forschungsgemeinschaft (DFG, German Research Foundation) under Germany’s Excellence Strategy – EXC 2002/1 “Science of Intelligence” – project number 390523135. M.R. was supported by the Deutsche Forschungsgemeinschaft (DFG; grants RO3579/8-1 and RO3579/10-1), as well as the European Research Council (ERC) under the European Union’s Horizon 2020 research and innovation program (grant agreement no. 865715).

